# Constitutive ghrelin receptor activity modulates AMPA receptor traffic and supports memory formation

**DOI:** 10.1101/2020.02.05.934463

**Authors:** Luís F. Ribeiro, Tatiana Catarino, Mário Carvalho, Sandra D. Santos, Luísa Cortes, Patricio O. Opazo, Lyn Rosenbrier Ribeiro, Daniel Choquet, José A. Esteban, João Peça, Ana Luísa Carvalho

**Affiliations:** CNC-Center for Neuroscience and Cell Biology, University of Coimbra, Coimbra, Portugal; MIT-Portugal Bioengineering Systems Doctoral Program; University of Bordeaux, Interdisciplinary Institute for Neuroscience, UMR 5297, Bordeaux, France; CNRS, UMR 5297, Bordeaux, France; Functional and Mechanistic Safety, Clinical Pharmacology and Safety Sciences, R&D AstraZeneca, Cambridge, CB2 0SL, UK; Bordeaux Imaging Center, UMS 3420, CNRS-Bordeaux University, US4 INSERM, Bordeaux, France; Department of Molecular Neurobiology, Centro de Biología Molecular “Severo Ochoa”, Consejo Superior de Investigaciones Científicas (CSIC) / Universidad Autónoma de Madrid, Madrid, Spain; Department of Life Sciences, University of Coimbra, Coimbra, Portugal

**Author notes:** These authors contributed equally to this work (Luís F. Ribeiro, Tatiana Catarino, Mário Carvalho). VIB Center for Brain and Disease Research, Leuven, Belgium; KU Leuven Department of Neurosciences, Leuven, Belgium. Clem Jones Centre for Ageing Dementia Research, Queensland Brain Institute, University of Queensland, St. Lucia, Queensland, Australia. Correspondence should be addressed to ALC.

**Keywords:** AMPA receptors, constitutive activity, ghrelin receptor, inverse agonism, memory

## Abstract

The ability of animals to store and retrieve food caches in the wild requires the integration of biological signals of hunger, satiety and memory. The role of ghrelin in regulating feeding and memory makes ghrelin receptors an important target to shape the required cellular and molecular responses. We investigated the effects of the high ligand-independent activity of the ghrelin receptor on the physiology of excitatory synapses. Blocking this type of activity produced a decrease in the synaptic content of AMPA receptors in hippocampal neurons and a reduction in GluA1 phosphorylation at Ser845. Impaired constitutive activity from the ghrelin receptor increased surface diffusion of AMPA receptors and impaired AMPA receptor synaptic delivery mediated by chemical long-term potentiation. These observations support a role for the constitutive activity of the ghrelin receptor in regulating AMPA receptor trafficking under basal conditions and synaptic plasticity. Accordingly, we found that blocking the ghrelin receptor constitutive activity impairs spatial and recognition memory.

**Impact statement:** This work uncovers a role for the constitutive activity of the ghrelin receptor in memory, and in the regulation of the synaptic levels of AMPA receptors, their mobility and synaptic plasticity. Underscoring the importance of deciphering the physiological role of constitutive ghrelin receptor activity, ghrelin receptor inverse agonism is now being considered as a therapy to treat alcohol use disorder.

## Introduction

Ghrelin is a peptide hormone believed to signal meal initiation (1, 2) and is found at the highest concentration in human plasma immediately before each meal (3). It is mainly secreted by X/A-like cells in the oxyntic glands of the stomach and intestine (4). Apart from effects on food intake (5) and feeding behavior (6, 7), ghrelin influences several other physiologic systems. For example, it is well established that ghrelin improves learning and memory (8, 9), regulates reward through its action on the mesolimbic dopamine system (10), modulates anxiety-(9) and depressive-like (11) behaviors, and affects long-term fear memory (12).

The actions of ghrelin are mediated by the growth hormone secretagogue receptor type 1a (GHSR-1a), a G protein-coupled receptor (GPCR), whose activation by ghrelin regulates gene expression, neuronal excitability and AMPA receptor trafficking (8, 13–15). In the brain, GHSR-1a is highly expressed in the hypothalamus, pituitary gland and hippocampus (16) and its expression levels increase during fasting (17–19). Interestingly, GHSR-1a displays unusually high constitutive activity, corresponding to approximately 50% of its maximal activity (20), that results from a natural shift in the equilibrium between its inactive and active conformations (21), in the absence of ligand. The ligand-independent GHSR-1a activity plays a role in the control of food intake and regulation of body weight (18, 19, 22–24), and in the acquisition of conditioned taste aversion (25). Human mutations that lead to a selective loss of constitutive activity of GHSR-1a, but that do not interfere with ghrelin-induced activation, are associated with familial short stature (26–28). The recently described liver-expressed antimicrobial peptide 2 (LEAP2) is an endogenous antagonist of the ghrelin receptor (29), which also exhibits inverse agonist activity, blocking ghrelin receptor constitutive activity (30, 31). LEAP2 plasmatic levels are lower in fasted states (29, 31), increase with body mass and blood glucose, and are higher in obesity (31), in a manner that is opposite to that of plasma acyl-ghrelin. These observations indicate that acyl-ghrelin and LEAP2 bidirectionally control ligand-dependent activity of the ghrelin receptor, and importantly that LEAP2 exerts endogenous control of the ligand-independent activity of the ghrelin receptor, which is physiologically relevant (32).

Ghrelin receptor knock-out animals display spatial and contextual memory impairments (33, 34), which can be attributed to the absence of ghrelin-triggered effects, but also to the loss of ligand-independent activity. However, the physiological importance of GHSR-1a constitutive activity for learning and memory has not been described. Here, using a combination of imaging, biochemical and electrophysiological approaches, and behavior analysis, we uncover a role for the constitutive activity of the ghrelin receptor in providing tonic control for the regulation of AMPA receptor traffic, influencing synaptic plasticity in the hippocampus and interfering with learning and memory *in vivo*. These finding should be taken into account given that inverse agonists of the ghrelin receptor are presently being tested in humans to treat alcohol use disorder (35, 36).

## Results

### AMPA-receptor surface and synaptic expression is regulated by ligand-independent activity of the ghrelin receptor

To test whether the constitutive activity of the ghrelin receptor could provide a tonic signal in the hippocampus and regulate AMPA receptor traffic, we evaluated whether inverse agonists of the ghrelin receptor affect the synaptic content of AMPA-type glutamate receptors in cultured hippocampal neurons. We used a newly described blood-brain barrier (BBB) permeable inverse agonist of the ghrelin receptor, AZ12861903 (AZ), which decreases the constitutive activity of the receptor (23), and also the more widely used [D-Arg^1^,D-Phe^5^,D-Trp^7,9^,Leu^11^]-substance P (SP-A), a well-established ghrelin receptor inverse agonist which does not cross the BBB (18, 20). Incubation of hippocampal neurons (15 DIV) with either of these compounds decreased the total cell surface levels of GluA1 (Figs. 1A,C and Supplementary Figs. 1A, B), and the levels of cell surface GluA1-containing AMPA receptors co-localized with the postsynaptic protein PSD95 and the presynaptic protein VGluT1 (Figs. 1A-D). In contrast, incubation with the ghrelin receptor antagonist JMV2959 did not significantly affect the total or synaptic levels of surface GluA1 (Supplementary Figs. 1C,D). Similarly to GluA1, GluA2 synaptic levels were decreased in neurons incubated with SP-A (Supplementary Figs. 1E,F). However, the incubation with SP-A did not affect synapse density in 15 DIV cultured hippocampal neurons (Supplementary Figs. 1E,G), measured by the colocalization of PSD95 and VGluT1 puncta. SP-A also decreased the total surface and synaptic levels of GluA1, as well as synapse density in older neurons (20 DIV, Supplementary Figs. 1H-J), but not in 7 DIV neurons (Supplementary Figs. 1K-M), which at this age present lower levels of expression of the ghrelin receptor (14).

**Figure 1.**
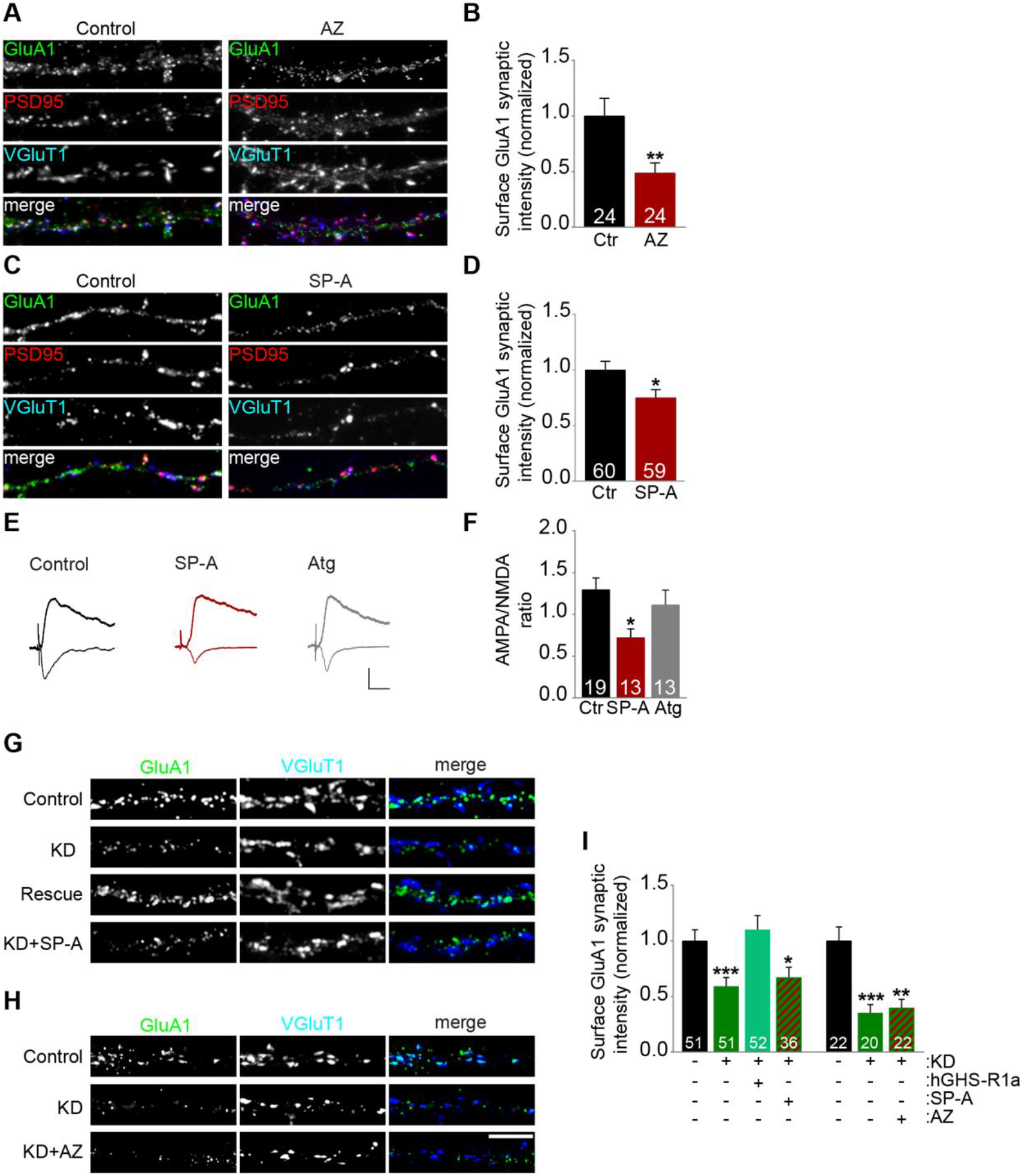
Ligand-independent activity of the ghrelin receptor controls synaptic levels of AMPAR and excitatory synaptic transmission in the hippocampus. (**A,C**) Hippocampal neurons [15 d *in vitro* (DIV)] were incubated with ghrelin receptor inverse agonists AZ12861903 (AZ) 50 nM (**A**) or [D-Arg^1^,D-Phe^5^,D-Trp^7,9^,Leu^11^]Substance P (SP-A) 1 μM (**C**) for 20 h and immunostained for surface GluA1 under non-permeabilizing conditions. After permeabilization, neurons were stained for MAP2, PSD95 and VGluT1 and analyzed for the total fluorescence intensity of GluA1 synaptic cluster (VGluT1/PSD95-colocalized) per synapse density. (**B**,**D**) Results are expressed as the percentage of control cells, and are averaged from 2–4 independent experiments. *\*P* < 0.05 and *\*\*P* < 0.01 by Mann-Whitney test. (**E**) Comparison of evoked synaptic AMPAR and NMDAR currents in 7 DIV organotypic hippocampal slices, in control condition, upon treatment for 20 h with SP-A, or with the ghrelin receptor antagonist (Atg; [D***-***Lys3]***-***GHRP-6, 100 μM). Scale bars: vertical, 50 pA; horizontal, 20 ms. (**F**) Average AMPA/NMDA ratios for SP-A-, antagonist-treated and control cells. *\*P* < 0.05 by Kruskal-Wallis test followed by Dunn’s post hoc test. (**G, H**) 15 DIV hippocampal neurons were transfected with constructs encoding luciferase shRNA-GFP (control), GHS-R1a shRNA-GFP (KD), GHS-R1a shRNA-GFP + hGHS-R1a (rescue) or GHS-R1a shRNA-GFP. Neurons expressing GHS-R1a shRNA-GFP were treated with SP-A **(G)** or AZ **(H)** for 20h. Neurons were immunostained for surface GluA1 under non-permeabilizing conditions. After permeabilization, neurons were stained for GFP, MAP2 and VGluT1, and analyzed for the total fluorescence intensity of GluA1 synaptic cluster (VGluT1-colocalized) per density of VGluT1 clusters. Scale bar represents 5 μm. (**I**) Results are expressed as the percentage of control cells, and are averaged from 3–4 independent experiments. *\*P* < 0.05 and *\*\*\*P* < 0.001 by Kruskal-Wallis test followed by Dunn’s post hoc test. All data are presented as mean ± SEM, and number in bars indicates number of cells analyzed for each condition. **See also Figure S1**.

To test whether the ligand-independent activity of the ghrelin receptor functionally modulates CA3-CA1 hippocampal excitatory transmission, organotypic hippocampal slices were treated with SP-A for 20h, and electrophysiological recordings were performed. The AMPA/NMDA ratio of synaptic responses decreased significantly after treatment with SP-A, compared with control neurons, whereas the ghrelin receptor antagonist [D-Lys^3^]-GHRP-6 did not affect AMPA/NMDA ratios (Figs. 1E, F), in agreement with low levels of ghrelin in the culture, and a specific role for the ghrelin receptor inverse agonist in inhibiting CA3-CA1 synaptic transmission. Together, these results suggest that the constitutive activity of GHSR regulates AMPA receptor levels and excitatory synaptic transmission under basal conditions.

We then silenced the expression of the ghrelin receptor in cultured hippocampal neurons using a short hairpin RNA (shRNA) containing a previously validated sequence (37) and tested for surface levels of GluA1. Knock-down of the ghrelin receptor decreased the levels of surface and synaptic GluA1, which were rescued when the shRNA was co-expressed with an shRNA-insensitive human form of the receptor (hGHS-R1a, Figs. 1G-I and Supplementary Fig. 1N). Notably, treatment with either SP-A or AZ did not affect the total surface or synaptic levels of GluA1 in neurons where the ghrelin receptor expression was silenced (Figs. 1G-I and Supplementary Fig. 1N), confirming the specificity of SP-A and AZ in targeting the ghrelin receptor.

### Ligand-independent activity regulates AMPA receptor surface mobility

To determine how the ligand-independent activity of the ghrelin receptor may regulate AMPA receptors, we tested for effects on the cell surface diffusion of GluA1 AMPA receptor subunit. We expressed superecliptic pHluorin (SEP)-GluA1 in cultured hippocampal neurons and took advantage of the single particle tracking approach to monitor individual AMPA receptor complexes (Figs. 2A-E). Hippocampal neurons were exposed to SP-A for 1 h and single nanoparticle imaging of SEP-GluA1 was performed thereafter. SP-A exposure significantly increased the surface diffusion of GluA1 (both the mean square displacement and diffusion coefficient were increased, Figs. 2A, B), decreased the fraction of synaptic immobile receptors (Fig. 2C) and decreased the synaptic residence time of GluA1-AMPA receptors (Fig. 2D).

**Figure 2.**
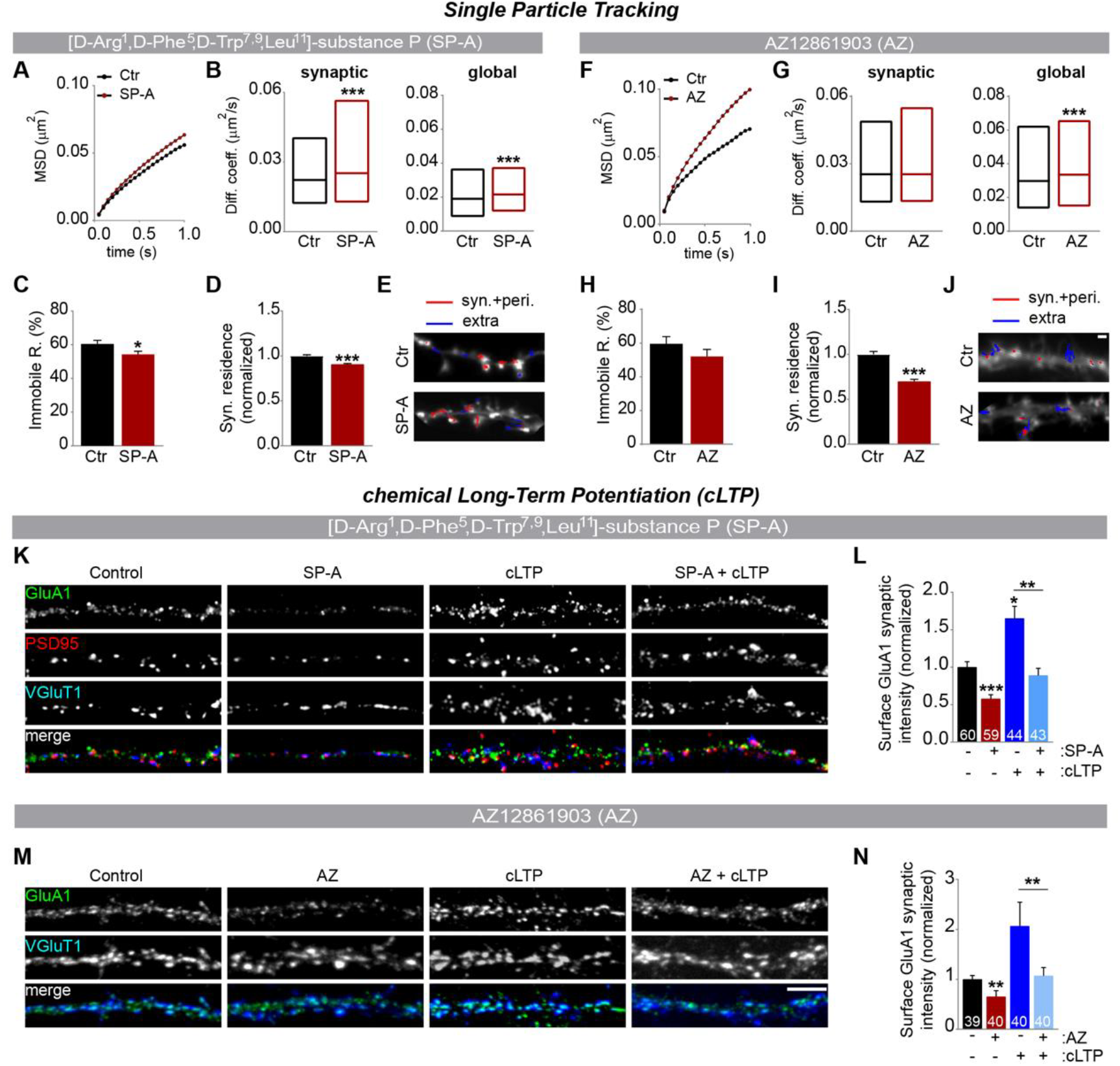
Blockage of the ligand-independent activity of the ghrelin receptor increases the cell surface diffusion of AMPAR, and prevents the activity-triggered delivery of AMPAR to synaptic sites. **(A–E)** Hippocampal neurons co-transfected with SEP-GluA1 and Homer1C-DsRed (11DIV) were incubated with SP-A 1 μM for 1 h (15DIV) prior to assessing GluA1 surface diffusion using quantum dots labelled antibodies for GFP to detect SEP-GluA1 (QD-GluA1). **(A)** GluA1 mean square displacement (MSD) versus time plots for control and SP-A-treated cells. **(B)** Surface diffusion coefficient of synaptic (left) and global (right) single QD-GluA1. Median diffusion (± 25%–75% IQR) of 8816–8607 trajectories. ***P < 0.001 by Mann-Whitney test. **(C)** Mean percentage (± SEM) of synaptic immobile GluA1-AMPAR in control and SP-A-treated cells. *P < 0.05 by Mann-Whitney test. **(D)** Mean synaptic residence time (± SEM) of GluA1-AMPAR in control and SP-A-treated cells. ***P < 0.001 by Mann-Whitney test. **(E)** Reconstructed GluA1 trajectories in the synaptic (red) and extrasynaptic compartments (blue). A minimum of 37 cells were analyzed in 3 independent experiments. **(F–J)** Hippocampal neurons transfected with Homer1C-DsRed (11 DIV) were incubated with AZ 50 nM for 1 h (15 DIV) prior to assessing GluA1 surface diffusion using quantum dots-labelled antibodies for GluA1 (QD-GluA1). **(F)** GluA1 mean square displacement (MSD) versus time plots for control and AZ-treated cells. **(G)** Surface diffusion coefficient of synaptic (left) and global (right) single QD-GluA1. Median diffusion (± 25%–75% IQR) of 1770–1303 trajectories. ***P < 0.001 by Mann-Whitney test. **(H)** Mean percentage (± SEM) of synaptic immobile GluA1-AMPAR in control and AZ-treated cells. **(I)** Mean synaptic residence time (± SEM) of GluA1-AMPAR in control and AZ-treated cells. ***P < 0.001 by Mann-Whitney test. **(J)** Reconstructed GluA1 trajectories in the synaptic (red) and extrasynaptic compartments (blue). 41 cells per condition were analyzed in 3 independent experiments. Scale bar represents 1 μm. **See also Figure S2**. (**K**) cLTP-induced insertion of synaptic GluA1. 19–20 DIV hippocampal neurons were either treated with SP-A for 20 h, submitted to cLTP, or pre-treated with SP-A and submitted to cLTP. Neurons were immunostained for surface GluA1 under non-permeabilizing conditions and analyzed for total fluorescence intensity of GluA1 synaptic clusters (VGLUT1/PSD95-colocalized) per synapse density. (**L**) Results are expressed as the mean percentage of control cells (± SEM), and are averaged from 3–4 independent experiments. Number in bars indicates number of cells analyzed for each condition. *\*P* < 0.05, *\*\*P* < 0.01 and *\*\*\*P* < 0.001 by Kruskal-Wallis test followed by Dunn’s post hoc test. (**M**) 19–20 DIV hippocampal neurons were either treated with AZ for 20 h, submitted to cLTP, or pre-treated with AZ and submitted to cLTP. Neurons were immunostained for surface GluA1 under non-permeabilizing conditions and analyzed for total fluorescence intensity of GluA1 synaptic clusters (VGluT1-colocalized) per synapse density. Scale bar represents 5 μm. (**N**) Results are expressed as the mean percentage of control cells (± SEM), and are averaged from 3–4 independent experiments. Number in bars indicates number of cells analyzed for each condition. *\*\*P* < 0.01 by Kruskal-Wallis test followed by Dunn’s post hoc test.

We repeated this experiment now applying AZ and using quantum-dots labeled antibodies against an extracellular region in GluA1, to follow endogenous AMPA receptors (Figs. 2F-J). Endogenous GluA1 also showed increased mean square displacement and diffusion coefficient and decreased synaptic residence time in neurons treated with AZ. Similarly, SP-A treatment increased the surface diffusion of endogenous GluA2 (Supplementary Figs. 2A-D). These data provide evidence that the constitutive activity of the ghrelin receptor contributes to decrease the surface diffusion of synaptic AMPA receptors, thereby increasing the synaptic content of AMPA receptors under basal conditions in hippocampal neurons.

### Effects of the constitutive activity of the ghrelin receptor on activity-induced synaptic incorporation of AMPA-receptor

We then assessed whether activity-induced synaptic incorporation of AMPA receptors is regulated by the ligand-independent activity of the ghrelin receptor. We used a neuronal culture model of chemical long-term potentiation (cLTP), in which activation of NMDA receptors triggers an increase in the expression of surface synaptic AMPA receptors (38). In agreement with previous reports, application of glycine, the co-agonist of NMDA receptors, in the absence of Mg^2+^ led to a significant increase in the synaptic expression of GluA1-AMPA receptors compared to control cells (Figs. 2K-N). However, this effect was blocked in neurons pre-incubated with either SP-A (Figs. 2K, L) or AZ (Figs. 2M, N), indicating that GHSR constitutive activity is necessary for AMPA receptors synaptic insertion upon cLTP.

### Intracellular signaling of ligand-independent activity of the ghrelin receptor

We tested cell signaling pathways downstream of the ghrelin receptor constitutive activity that could result in altered trafficking of AMPA receptors. We found that upon blockade of the ligand-independent activity of the ghrelin receptors in organotypic hippocampal slices there was a decrease in the phosphorylation of GluA1 at Ser845 (Fig. 3A), a PKA phosphorylation site critical for priming AMPA receptors for synaptic insertion (39). Additionally, a decrease in the phosphorylation of CaMKIV (Fig. 3B) was detected, whereas no changes were found in the phosphorylation of GluA1 at Ser831, in the phosphorylation of stargazin or in the phosphorylation of Akt (Figs. 3C-E). These results suggest that PKA activation downstream of the constitutive activity of the ghrelin receptor may result in phosphorylation of GluA1 at Ser845 and contribute to maintaining a population of AMPA receptors available for synaptic insertion.

**Figure 3.**
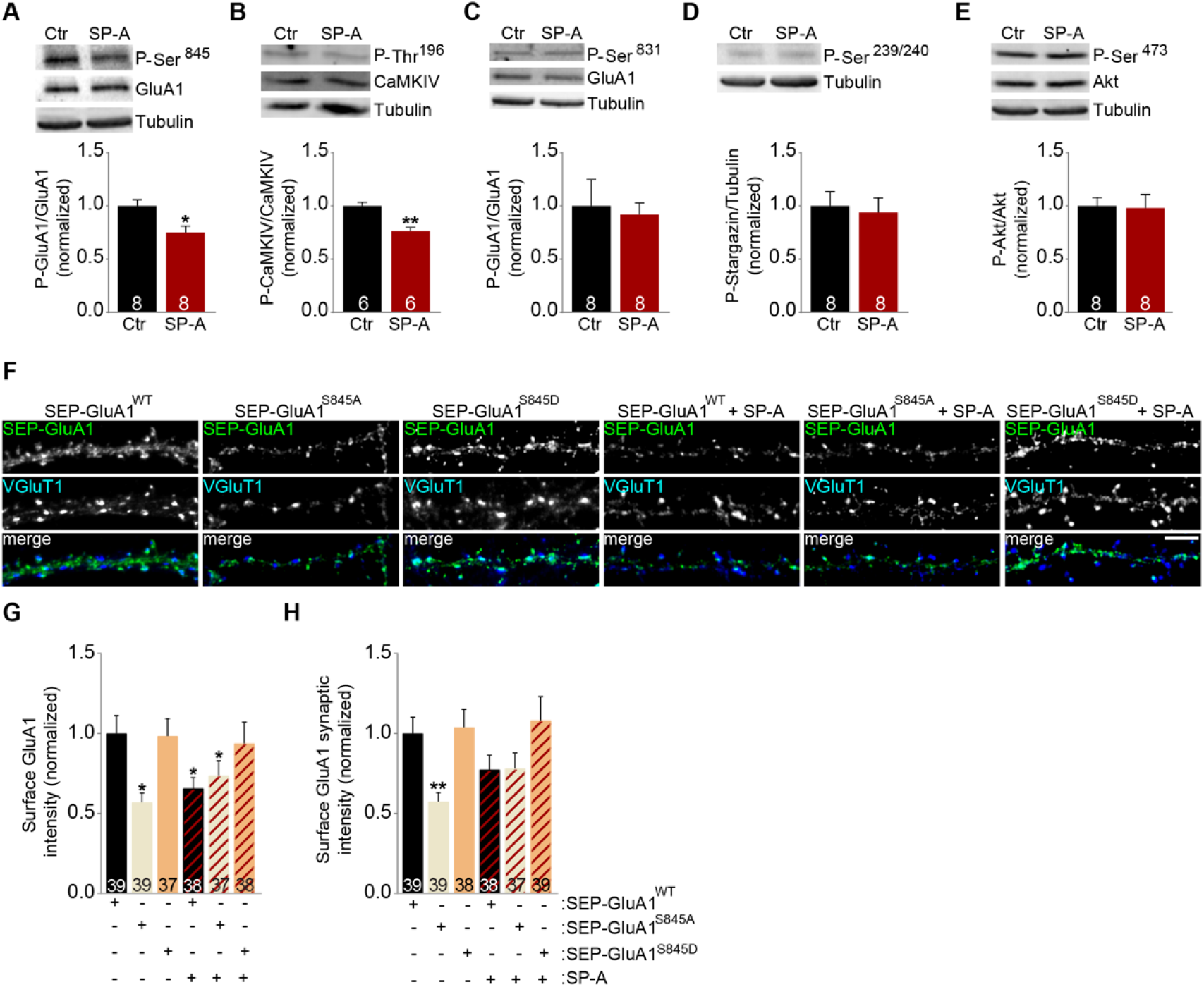
Signaling pathways downstream of the ghrelin receptor constitutive activity. (**A-E**) Western blot analysis of protein extracts from 7 DIV organotypic hippocampal slices incubated with culture medium or medium containing SP-A (1 μM) for 20 h. Primary antibodies detected phosphorylation of GluA1 at Ser^845^ (**A**), phosphorylarion of CaMKIV at Thr^196^ (**B**), phosphorylation of GluA1 at Ser^831^ (**C**), phosphorylation of stargazin at Ser^239/240^ (**D**), phosphorylarion of Akt at Ser^473^ (**E**), a residue targeted by a PI3 kinase downstream signaling pathway. Total GluA1 (**A, C**), total CaMKIV (**B**) and total Akt (**E**) were also detected. Tubulin was used as a loading control in all cases. The graphs represent the quantification of band intensities relative to control extracts. Error bars represent SEM. The statistical significance was calculated using the Mann-Whitney test (\**P* < 0.05 and \*\**P* < 0.01). Number in bars indicates number of independent experiments. (**F-H**) 15 DIV hippocampal neurons expressing SEP-GluA1 (control), SEP-GluA1S845A or SEP-GluA1S845D, non-treated or treated with SP-A for 20 h, were immunostained for surface GFP under non-permeabilizing conditions. After permeabilization, neurons were labeled with Alexa568-conjugated secondary antibody to reveal surface GluA1. Neurons were also stained for MAP2 and VGluT1, and analyzed for the cell-surface fluorescence intensity of GFP clusters, total and synaptic (VGluT1-colocalized), per density of VGluT1 clusters. Scale bar represents 5 μm. (**G,H**) Results are expressed as the percentage of control cells, and are averaged from 3 independent experiments. *\*P* < 0.05 and *\*\*P* < 0.01 by Kruskal-Wallis test followed by Dunn’s post hoc test. All data are presented as mean ± SEM, and number in bars indicates number of cells analyzed for each condition.

To test whether the ghrelin receptor constitutive activity contributes to AMPA receptor traffic through effects on GluA1 phosphorylation at Ser845, we evaluated whether SP-A affects the cell surface and synaptic levels of phosphodead and phosphomimetic mutants of GluA1 at Ser 845 (Ser845A and Ser845D, respectively). We found that contrarily to wild-type GluA1, SP-A treatment did not alter the cell surface or synaptic levels of either of these mutants (Figs. 3F-H), suggesting GluA1 phosphorylation at Ser845 as one mechanism through which the constitutive activity of the ghrelin receptor regulates AMPA receptor traffic.

### Constitutive activity of the ghrelin receptor is relevant for memory formation

Given the role of the hippocampus and excitatory transmission in spatial memory (40), and that ghrelin receptor knock-out mice present memory impairments (33, 34), we tested whether ghrelin receptor constitutive activity plays a role in memory formation by evaluating performance in the novel object recognition test (41) in mice injected with the BBB-permeable inverse agonist of the ghrelin receptor AZ. During the familiarization session, animals were allowed to explore two identical objects for 10 min. After 6 h, one of the objects was replaced with a novel object, and the percentage of time exploring either object was measured (test session, Fig. 4A). Whereas control animals explored the novel object a higher number of times and for longer, animals treated with the ghrelin receptor inverse agonist prior to the familiarization session did not show a preference for either object, as measured by the number of explorations of each object (Fig. 4B) or time spent with each object (Fig. 4C). To account for changes in motor behavior, before and after drug administration, we also measured total distance travelled by animals in the arena and did not find significant alterations between vehicle- and drug-treated animals (Supplementary Fig. S3A).

**Figure 4.**
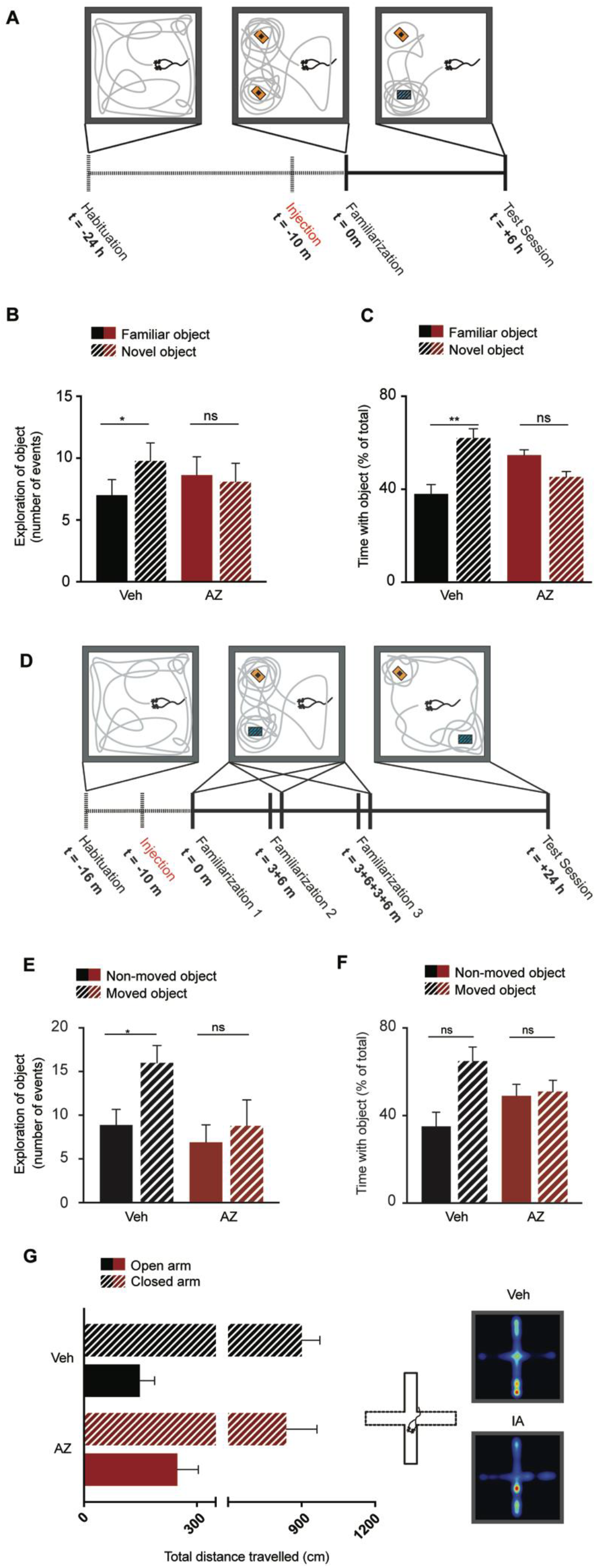
Administration of an inverse agonist of the ghrelin receptor impairs recognition and spatial memory. (**A**–**F**) Male C57/BL6 mice were divided into two groups, vehicle (Veh)-treated animals and animals receiving 100 mg/kg of AZ. **(A-C)** Mice were habituated to the arena for 10 min without any objects and, after a delay of 24 h, were injected i.p. with either vehicle or 100 mg/kg AZ, before being re-introduced for 10 min in the arena now containing two identical objects. Following a delay of 6 h, animals were re-introduced in the arena where a novel object replaced one of the familiar objects, and animals explored the arena for 10 min. Only the first 5 min of the test were considered. (**A**) Schematic representation of a novel object recognition test with “habituation”, “drug injection”, “familiarization” and “test session” time line. (**B**) Number of interactions with objects during test session is expressed as a percentage (± SEM) of all interactions with objects, for vehicle- and AZ-treated groups; n=13 Veh, n=11 AZ. (**C**) Time spent with objects during the test session is expressed as a percentage (± SEM) of total duration of interaction with both objects, for vehicle- and AZ-treated groups. **(D-F)** A new cohort of mice were habituated to the arena for 6 min in the absence of objects and were injected with either vehicle or 100 mg/kg AZ immediately after. Injected animals were returned to the homecage for 10 min before returning to the arena where two different objects were placed in a specific location. Mice were allowed to explore the objects for 6 mins (familiarization), and this session was repeated 3 times with 3 min interval between each repetition. After 24 h, the animals were returned to the arena, where one of the objects was placed in a new location, and were allowed to explore for 6 min. **(D)** Schematic representation of an object displacement recognition test with “habituation”, “drug injection”, “familiarization” and “test session” time line. **(E)** Number of interactions with objects during test session is expressed as a percentage (± SEM) of all interactions with objects, for vehicle- and AZ-treated groups; n=8 Veh, n=9 AZ. **(F)** Time spent with objects during the test session is expressed as a percentage (± SEM) of total duration of interaction with both objects, for vehicle- and AZ-treated groups. **(B,C,E,F)** Comparisons between groups were performed using the Wilcoxon matched-pairs signed rank test; * P<0.05, **P < 0.01. (**G**) Total distance travelled in an elevated plus maze was assessed to determine anxiety-like behavior following administration of AZ at 100 mg/kg (n=5) when compared to vehicle injections (n=6). No significant differences were found in distance travelled in the open arms of the elevated plus maze, Mann-Whitney test P > 0.05. **See also Figure S3**.

To test whether the constitutive activity of the ghrelin receptor is relevant for spatial memory, we used the object displacement test (42) (Fig. 4D). During the habituation session the animals explored an open-field arena in the absence of objects for 6 min; immediately after, the animals were treated with ghrelin receptor inverse agonist or vehicle injection. The animals returned to the open-field 10 min post-injection, and 2 different objects were present in specific locations. This familiarization session lasted 6 min and was repeated twice. After 24h, the animals were tested in the open-field with one of the objects displaced to a different location. Our results show that animals injected with vehicle preferentially explored the moved object, whereas animals injected with the inverse agonist did not show such preference (Figs. 4E,F). Total distance travelled was not affected by injection of the ghrelin receptor agonist (Supplementary Fig. 3B). Since anxiolytic effects have been observed in mice administered with ghrelin (11), we tested performance in the elevated plus maze after blocking constitutive activity of the ghrelin receptor. Consistent with prior results using knock-out mice for the ghrelin receptor (11), blockade of the constitutive activity of the ghrelin receptor did not affect performance in the elevated plus maze (Fig. 4G and Supplementary Figs. S3C,D).

Our results indicate that acute blockade of GHSR-1a constitutive activity impairs performance during the novel object recognition task and in the object displacement recognition task, which suggests that tonic activity of the ghrelin receptor is important for learning and memory.

## Discussion

The constitutive activity of the ghrelin receptor has been previously described to regulate food intake and body weight (18, 19), but its involvement in the regulation of hippocampal synaptic transmission and plasticity, memory and learning has not been directly addressed. Here, we provide strong evidence that the ligand-independent activity of the ghrelin receptor regulates AMPA receptors at the synapse and the formation of spatial memories.

We first tested whether two inverse agonists of the ghrelin receptor affect the synaptic content of AMPA receptors, which are critical for synaptic plasticity processes thought to underlie learning and memory (43). We observed that the GHSR constitutive activity regulates the levels of synaptic GluA1 and GluA2 AMPA receptors in cultured hippocampal neurons, and the AMPA/NMDA ratio in hippocampal organotypic slice cultures. In agreement with these observations, we found a decrease on the synaptic levels of GluA1 upon silencing of the expression of the ghrelin receptor. We also confirmed the specificity of both inverse agonists of the ghrelin receptor, since they showed no effects on GluA1 in neurons depleted of the ghrelin receptor expression.

We further explored how the GHSR constitutive activity impacts the synaptic and surface levels of AMPA-receptors, which are controlled both by exocytic/endocytic receptor traffic and by the recruitment to the synapse of extrasynaptic receptors preexisting at the neuronal plasma membrane (44). Using single particle imaging, we evaluated whether the ghrelin receptor ligand-independent activity affects the cell surface diffusion of AMPA receptors, which is critically important for synaptic plasticity underlying learning and memory (45). We observed that blocking ghrelin receptor constitutive activity enhanced the mobility of synaptic GluA1 and GluA2 AMPA-receptors, and decreased the residence of GluA1 and GluA2 at synapses.

Ligand-independent activity of the ghrelin receptor may be required for synaptic plasticity processes. Indeed, using a neuronal culture model of cLTP we observed that both inverse agonists of the ghrelin receptor block LTP, as measured by the synaptic accumulation of GluA1. These observations suggest that the high constitutive signaling of the ghrelin receptor generates a signaling set-point to regulate the levels of synaptic AMPA receptors and synaptic plasticity in the hippocampus. Our data further indicate that this control is produced through the phosphorylation of GluA1 at serine 845. Phosphorylation of GluA1 serine 845 regulates extrasynaptic membrane trafficking of GluA1, and primes AMPA receptors for synaptic insertion upon the induction of synaptic plasticity (39). Interestingly, phosphorylation of this site is also required for retention of spatial learning (46).

In order to assess the physiologic relevance of the ligand-independent activity of the ghrelin receptor we tested the effect of its blockade on the memory for object novelty and location. We observed that blocking ligand-independent activity of the ghrelin receptor impairs memory assessed by the novel object recognition and object displacement recognition tests. This effect is in agreement with the observation that ghrelin receptor knockout mice perform poorly in memory tests (33, 34), and suggests a clear role in memory for the unusually high constitutive activity of the ghrelin receptor, an intrinsic feature of this receptor (47).

Besides signaling in response to ghrelin, and in the absence of the ligand, the ghrelin receptor has recently been shown to modulate dopamine signaling through heterodimerization with dopamine receptors DRD1 and DRD2 (48, 49). In the hippocampus, GHSR-1a and DRD1 form heteromers that are activated by DRD1 agonists to induce intracellular Ca^2+^ mobilization, activation of early synaptic plasticity markers, and to modulate memory (49). Importantly, the dopamine-induced effect on Ca^2+^ signaling is independent of the constitutive activity of the ghrelin receptor in the GHSR-1a:DRD1 complex (49). This suggests that the role of the constitutive activity of the ghrelin receptor on memory here described runs parallel to the effects of dopamine on memory through the GHSR-1a:DRD1 complex.

The GHSR-1a constitutive activity has also been shown to reduce presynaptic Ca_v_2 currents and GABA release in hypothalamic and hippocampal neurons (50, 51), by reducing the cell surface expression of Ca_v_2 channels (52). Our results complement this observation, but work should be done to explore how the effects of the constitutive activity of the ghrelin receptor on the inhibitory and excitatory systems contribute to memory formation.

It was recently reported that the melanocortin receptor accessory protein 2 (MRAP2) controls GHSR-1a signaling by inhibiting its constitutive activity, as well as by increasing its G protein-mediated signaling and blocking the recruitment and signaling of β-arrestin elicited by ghrelin binding (53). Disruption of the gene for MRAP2 has been associated with obesity in animal models and humans (54). MRAP2 mRNA has low expression in the hippocampus of both animals (54, 55) and humans (56), which suggests that constitutive activity of the ghrelin receptor is unimpeded in this region and is thus more likely to have an influence in hippocampal excitatory synapse protein dynamics and hippocampus-dependent behavior.

Recent evidence suggests that the constitutive activity of the receptor is endogenously regulated by the plasmatic levels of LEAP2 (49), which are proportional to the levels of adiposity and blood glucose (31), and also potentially by the levels of expression of the ghrelin receptor (18), which in turn change according to the animal’s feeding status (17, 18). Therefore, our observations support a physiological mechanism in which the internal metabolic state of animals exerts control over cognitive processes. Finally, the role of the ghrelin receptor constitutive activity reported in this work should be taken into account when considering its inverse agonists as anti-obesity agents (57), or their use for the treatment of alcohol use disorder (35, 36).

## Materials and Methods

### Animals and Behavior

For the behavior experiments, 8-15 weeks old male C56BL/6 mice were kept in the Animal Facility of the CNC/Faculty of Medicine of the University of Coimbra with access to food and water *ad libitum*. The environment was kept in temperature and humidity controlled conditions under a 12h dark-light cycle (light period 6h00-18h00). Behavioral testing was reviewed and approved by the animal use and ethics committee (ORBEA) of the CNC/Faculty of Medicine, University of Coimbra, and by the Portuguese national authority for animal experimentation (DGAV), and all procedures were performed according to the guidelines of the DGAV and Directive 2010/63/EU of the European Parliament.

The novel object recognition task was adapted from Leger and colleagues (41), the object displacement test was adapted from Oliveira and colleagues (42) and the elevated plus maze was performed according to (58). These tests are described in more detail in SI Materials and Methods.

### Immunocytochemistry

For labeling surface GluA1-containing AMPA receptors, live neurons were incubated for 10 min at room temperature using an antibody against an extracellular epitope in the GluA1 N-terminus diluted in conditioned neuronal culture medium or extracellular solution (used for chemical LTP). Neurons were then fixed and stained as previously described (59). For labeling GluA2-containing AMPA receptors, neurons were fixed and then incubated overnight with an anti-GluA2 antibody diluted 1:100 in 3%BSA/PBS, at 4°C. Neurons were then stained as previously described (59).

### Quantitative imaging analysis

Imaging was performed on a Zeiss Axio Observer Z1 microscope using a Plan Apochromat 63×/1.4 NA oil objective, and an AxioCam HRm CCD camera. Images were quantified using image analysis software (ImageJ). For quantification, sets of cells were cultured and stained simultaneously, and imaged using identical settings. The region of interest was randomly selected avoiding primary dendrites, and dendritic length was measured using MAP2 staining. Measurements were performed in 2–5 independent preparations, and at least 7 cells per condition were analyzed for each preparation. Quantitative imaging quantification was performed as previously described (59).

### Statistical analysis

Statistical differences were calculated according to non-parametric tests in most cases; Mann-Whitney test was used to compare statistical differences between any two groups. Comparisons between multiple groups were performed with the Kruskal-Wallis analysis of variance followed by Dunn’s multiple Comparison test or with 2-way ANOVA with Bonferroni test for correction of multiple comparisons.

Additional details are available in SI Materials and Methods.

## Acknowledgments

The authors thank AstraZeneca for sharing AZ12861903 through the AstraZeneca Open Innovation program. J.P. would like to acknowledge the support of the FCT IF Programme (IF/00812/2012). This work was supported by a NARSAD Independent Investigator Grant from the Brain and Behavior Research Foundation, by national funds through the Portuguese Science and Technology Foundation (FCT; POCI-01-0145-FEDER-007440 and POCI-01-0145-FEDER-PTDC/SAU-NMC/4888/2014 and POCI-01-0145-FEDER-28541), and by the European Regional Development Fund (ERDF), through the Centro 2020 Regional Operational Programme under project CENTRO-01-0145-FEDER-000008:BrainHealth 2020.

## Author Contributions

LFR, TC, MC, SDS, LC, PO and LRR performed experiments. LRR provided new reagents. LFR, TC, MC, SDS, LC, PO and LRR analyzed data. LFR, DC, JAE, JP and ALC designed experiments. LFR, MC, JP and ALC wrote the paper.

## Supplemental Information

**Supplementary Figure S1.**
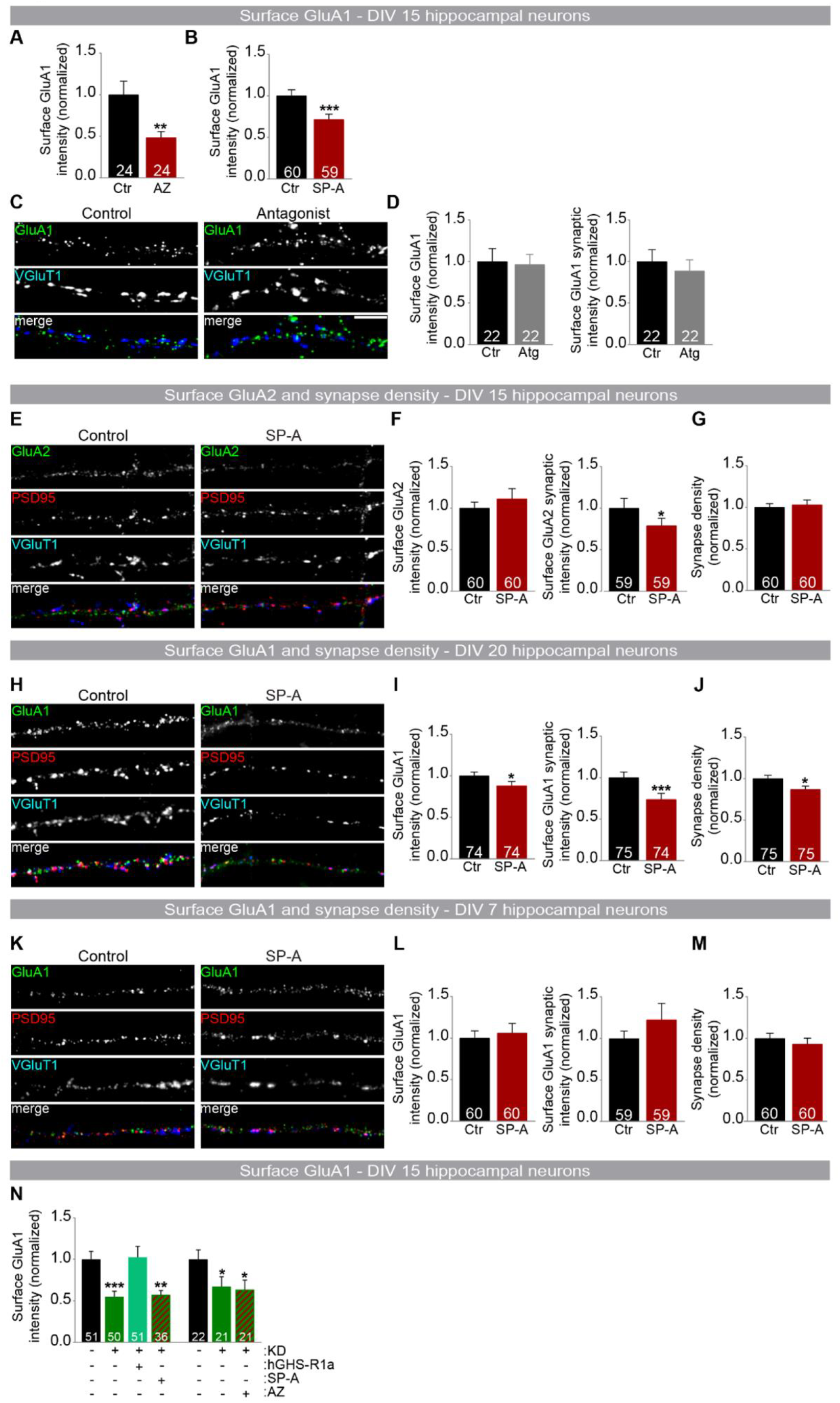
(Related to Figure 1) (**A,B**) Hippocampal neurons [15 d *in vitro* (DIV)] were incubated with ghrelin receptor inverse agonists [D-Arg^1^,D-Phe^5^,D-Trp^7,9^,Leu^11^]-Substance P [SP-A] 1 μM (**A**) or AZ12861903 [AZ] 50 nM (**B**) for 20 h, and immunostained for surface GluA1 under non-permeabilizing conditions. After permeabilization, neurons were stained for MAP2, PSD95 and VGluT1 and analyzed for the total fluorescence intensity of GluA1 cell-surface puncta normalized to synapse density. Results are expressed as the percentage of control cells, and are averaged from 2–4 independent experiments. *\*\*P* < 0.01 and *\*\*\*P* < 0.001 by Mann-Whitney test. (**C,D**) Hippocampal neurons [15 d *in vitro* (DIV)] were incubated with the ghrelin receptor antagonist JMV2959 100 μM for 20 hours and immunostained for surface GluA1 under non-permeabilizing conditions. After permeabilization, neurons were stained for MAP2 and VGluT1 and analyzed for the total fluorescence intensity of GluA1 synaptic cluster (VGluT1-colocalized) per synapse density. Results are expressed as the percentage of control cells, and are averaged from 2 independent experiments. (**E-M**) Hippocampal neurons 15 DIV (**E**), 20 DIV (**H**) and 7 DIV (**K**) were incubated with the ghrelin receptor inverse agonist SP-A 1 μM for 20 h and immunostained for surface GluA2 (**E**) or GluA1 (**H**,**K**) under non-permeabilizing conditions. After permeabilization, neurons were stained for MAP2, PSD95 and VGluT1 and analyzed for the total fluorescence intensity of GluA2 or GluA1 cell-surface puncta and total fluorescence intensity of GluA2 or GluA1 synaptic clusters (VGluT1/PSD95-colocalized) per synapse density (**F,I,L**). Neurons were also analyzed for synapse density (VGluT1 puncta positive for PSD95) (**G,J,M**). Scale bar represents 5 μm. Results are expressed as % of control cells, and are averaged from 4–5 independent experiments. Error bars represent SEM. *\*P* < 0.05 and *\*\*\*P* < 0.001 by Mann-Whitney test. Number in bars indicates number of cells analyzed for each condition. (**N**) Hippocampal neurons transfected at 7 DIV with Luciferase shRNA-GFP (control), GHS-R1a shRNA-GFP (KD), GHS-R1a shRNA-GFP + hGHS-R1a (rescue) or with GHS-R1a shRNA-GFP and treated at 15 DIV with SP-A or AZ were analyzed for the total fluorescence intensity of GluA1 cell-surface puncta normalized to synapse density. Results are expressed as % of control and are averaged from 3–4 independent experiments. Error bars represent SEM. Number in bars indicates number of cells analyzed for each condition. The statistical significance was calculated using the Kruskal-Wallis test followed by the Dunn’s Multiple Comparison test (\*\**P* < 0.01 and \*\*\**P*< 0.001).

**Supplementary Figure S2.**
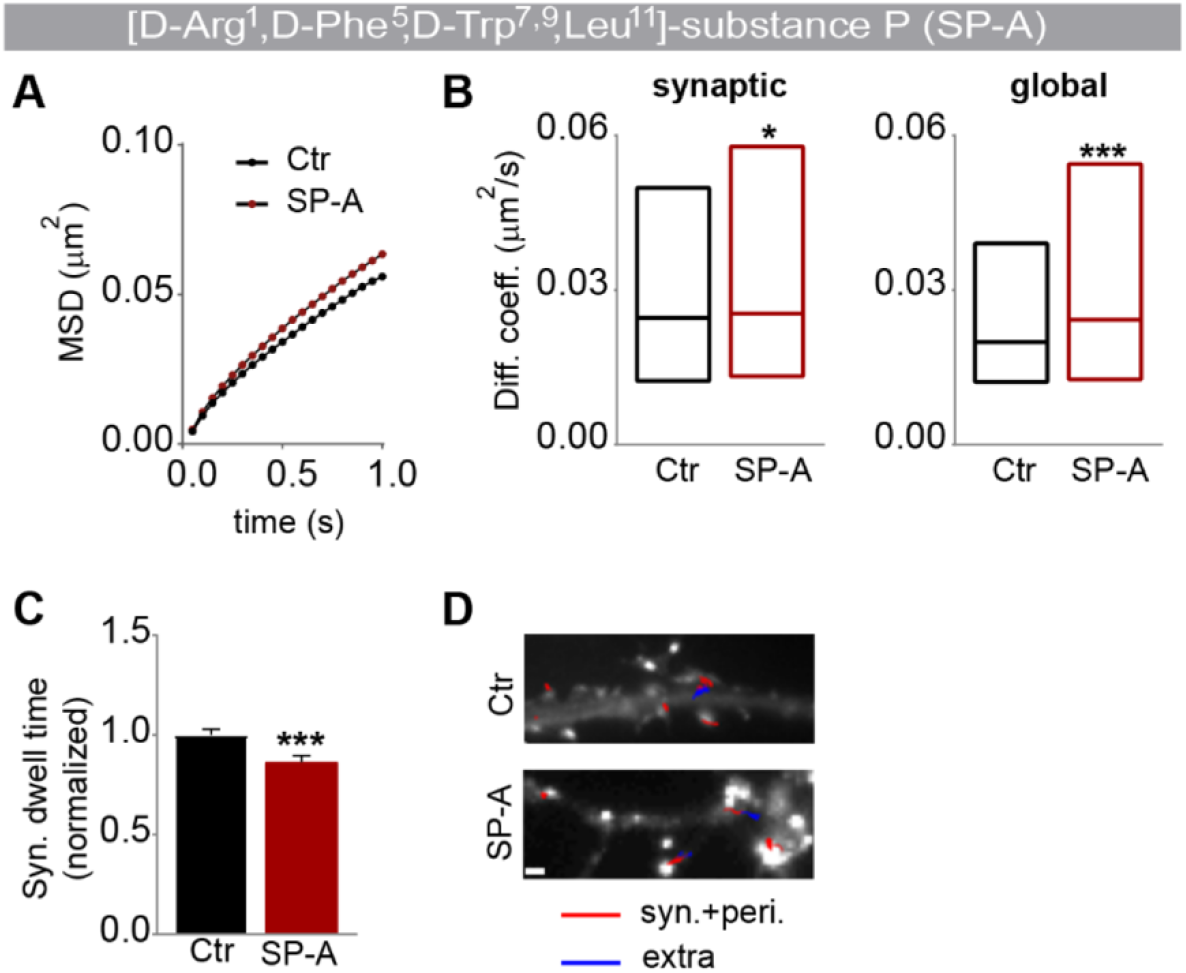
(Related to Figure 2) (**A**–**D**) Hippocampal neurons transfected with Homer1C-GFP (11 DIV) were incubated with SP-A 1 μM for 1 h (15 DIV) prior to assessing GluA2 surface diffusion using quantum dots labelled antibodies for GluA2 (QD-GluA2). (**A**) GluA2 mean square displacement (MSD) versus time plots for control and SP-A-treated cells. (**B**) Surface diffusion coefficient of synaptic (left) and global (right) single QD-GluA2. Median diffusion (± 25%– 75% IQR) of 4356-2288 trajectories. *\*P* < 0.05 and *\*\*\*P* < 0.001 by Mann-Whitney test. (**C**) Synaptic dwell time (± SEM) of GluA2 in control and SP-A-treated cells. *\*\*\*P* < 0.001 by Mann-Whitney test. (**D)** Reconstructed GluA2 trajectories in the synaptic (red) and extrasynaptic compartments (blue); scale bar represents 1 μm. A minimum of 33 cells were analyzed in 3 independent experiments.

**Supplementary Figure S3.**
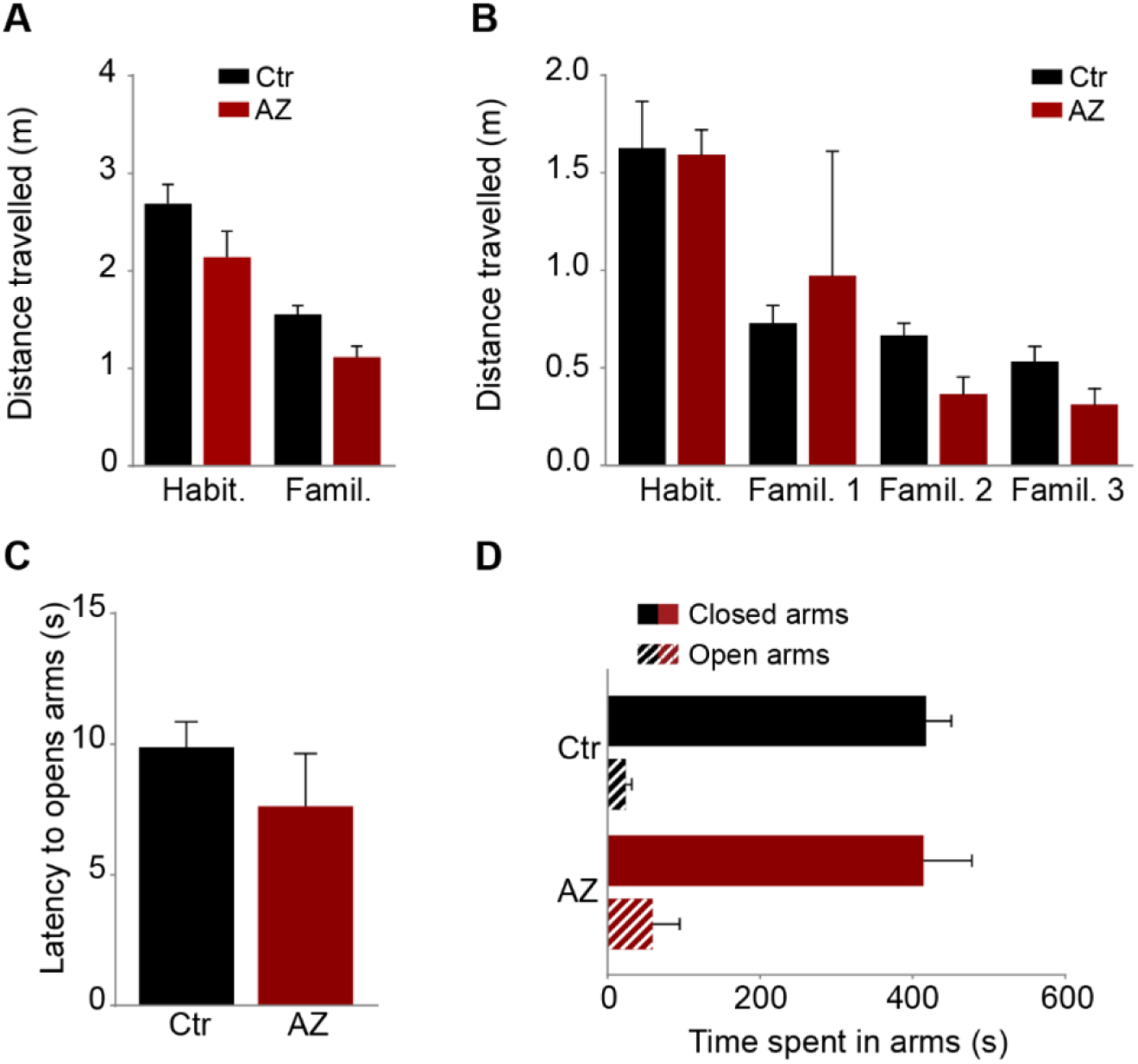
(Related to Figure 4) **(A)** Distance travelled (± SEM) by each animal during the first 5 min of novel object recognition test sessions: “habituation” and “familiarization”; n=13 vehicle (Ctr), n=11 AZ. **(B)** Distance travelled (± SEM) by each animal during the first 5 min of the object displacement recognition test sessions: “habituation” and “familiarization”; n=8 vehicle (Ctr), n=9 AZ. Comparison between groups was performed using 2 way ANOVA with Bonferroni test for correction of multiple comparisons. (**C**) Latency to enter open arms (± SEM) and (**D**) time spent on each arm of the elevated plus maze (± SEM); n=6 Ctr, n=5 AZ. No significant differences were found for either measurements between groups, using Mann-Whitney test.

## Supplemental Materials and Methods

### Behavior

The novel object recognition task was adapted from Leger and colleagues (1). This task consisted of 3 phases: In a first phase the animals freely explored the empty open-field arena for 10 min (habituation phase). Twenty four hours after, the animals were allowed to explore two similar, symmetrically disposed objects, for 10 min. Ten minutes before this phase the animals were submitted to intraperitoneal injection of either the drug or the vehicle, and stayed in an empty transport cage before entering the training phase. Six hours after the training phase, the animals were exposed to 2 objects located in the same positions as previously, but this time one of the objects was substituted by a new object that the animal had not contacted previously (test phase).

The object displacement test was adapted from Oliveira and colleagues (2). The test took place during 2 days. During the first day the animals were allowed to explore an empty open-field for 6 min (habituation phase). Immediately after, the animals were intraperitoneally injected with the inverse agonist of the ghrelin receptor or its respective vehicle and placed in their homecage. After 10 minutes, the animals explored two different objects placed in a specific location of the open field for 6 minutes (training phase). The animals were then returned to their homecage and waited for 3 minutes. Two more similar training phases were conducted, with a 3 minute waiting period in between. On the next day, the animals returned to the open field, where one of the objects was placed in a new location, and were allowed to explore the objects for 6 minutes (test phase).

The objects and their positions were randomized for both tests. The used objects correspond to the described by Leger and colleagues (1). The arena and the objects were carefully cleaned before running each animal and in between phases. The test was conducted at a room temperature of 23°C and 15 lux at the center of the arena (homogenously distributed light). Videos of the test were acquired using Noldus Ethovision software and scoring was performed blinded to treatment of the animals, using Noldus Observer.

Elevated plus maze was performed with a maze made in-house according to previously described specifications (3), and using an adapted protocol (4). Animals were weighted and injected with the corresponding dose of inverse agonist or vehicle 10 min before starting the test, and stayed in an empty transport cage. The test started by putting the animals in the central part of the maze with the nose aligned with the closed arms, and run for 10 min. The test was conducted under 100 lux at the center of the arena. The arena was carefully cleaned before and after each run. Videos of the test were acquired and automatically quantified using Noldus Ethovision.

### Materials

The GHS-R1a inverse agonist [D-Arg^1^,D-Phe^5^,D-Trp^7,9^,Leu^11^]-substance P (SP-A) was purchased from Bachem (Bubendorf, Switzerland), the AZ12861903 (AZ) GHS-R1a inverse agonist was kindly provided by AstraZeneca. The GHS-R1a agonist MK-0677 was purchased from Axon Medchem (Groningen, The Netherlands). The GHS-R1a antagonist [D-Lys^3^]-GHRP-6, TTX, and picrotoxin were purchased from Tocris Bioscience (Bristol, UK). The anti-Tubulin antibody was purchased from Sigma-Aldrich (Sintra, Portugal); the anti-Akt, anti-P-Ser473(Akt), anti-PSD95 (rabbit), and anti-CaMKIV antibodies were obtained from Cell Signaling (Danvers, MA, USA); the anti-GluA1, anti-GluA2, anti-P-Ser845(GluA1), anti-P-Ser239/240 (Stargazin) and anti-VGluT1 antibodies were from Millipore (Madrid, Spain); the anti-MAP2 antibody was from Abcam (Cambridge, UK), the anti-P-S831(GluA1) antibody was from Tocris Bioscience (Bristol, UK), the anti-PSD95 (mouse) antibody was from Affinity BioReagents (Golden, USA), and the anti-P-Thr196 (CaMKIV) antibody was from Santa Cruz Biotechnology Inc (Santa Cruz, CA); the anti-GFP (rabbit) antibody was from MBL International (Woburn, USA) and the anti-GFP (mouse) antibody was from Roche (Amadora, Portugal). Quantum dots (QDs) 655 Goat F(ab’)2 anti-mouse IgG conjugate (H^+^L) were purchased from Invitrogen (Barcelona, Spain). The antibody for the N-terminus of GluA1 was a kind gift from Dr. Andrew Irving (University College Dublin). All other reagents were purchased from Sigma-Aldrich (Sintra, Portugal), Fisher Chemicals or from Merck (Darmstadt, Germany) unless specified otherwise.

### DNA constructs

The GFP-tagged GHS-R1a construct (5) was a kind gift from Dr. Helen Wise (The Chinese University of Hong Kong) and SEP-GluA1 was generously provided from Helmut Kessels (Netherlands Institute for Neuroscience). For the generation of the short hairpin interfering RNA construct targeting the ghrelin receptor, a previously described and validated sequence (6) was used. Complementary oligonucleotides, each containing a unique 19-nt sequence derived from within the target mRNA transcripts of *ghsr1a* gene (NM_032075) targeting nucleotides 79-96 (GACTCACTGCCTGACGAAC) (6), were annealed and subcloned into the HpaI/XhoI sites of the U6 promoter-driven short hairpin RNA expression vector pLentiLox3.7(CMV)EGFP, which co-expresses EGFP under the CMV promoter. The control shRNA that targets firefly luciferase was described previously (7). Homer1C-DsRed and Homer1C-GFP were previously described (8).

### Application of GHS-R1a agonists and antagonists

Hippocampal organotypic slices (6 DIV) were treated with the GHS-R1a inverse agonist [D-Arg^1^, D-Phe^5^,D-Trp^7,9^,Leu^11^]-substance P (SP-A, 1μM) for 20 h, or chronically treated with the GHS-R1a antagonist [D-Lys^3^]-GHRP-6 (100 μM) from 3 DIV up to 7 DIV. Hippocampal neurons in culture were incubated with the GHS-R1a inverse agonists (SP-A and AZ) and antagonist JMV2959. The compounds were added directly to the culture medium. AZ for injection *in vivo* was dissolved in 95% beta-hydroxypropylcyclodextrin (**β**-hpC)/5% v/v DMSO. **β**-hpC was prepared at 25 % w/v in Sorenson’s buffer pH 5.5. All the injected solutions were prepared in sterile conditions. The drug and vehicle were injected intraperitoneally at volumes of 100-150 µl. The dose of 100 mg/kg was based on previously described doses by McCoull and colleagues (9).

### Neuronal and slice cultures

Primary cultures of rat hippocampal neurons were prepared as previously described (10). Hippocampal slices were prepared from young Wistar rats of either sex (postnatal day 5–6) as previously described (11).

### Neuron transfection

DNA constructs [Luciferase shRNA-GFP, GHS-R1a shRNA-GFP (Knock-down), hGHS-R1a (rescue), SEP-GluA1, Homer1C-DsRed and Homer1C-GFP] were expressed in primary cultures of hippocampal neurons at 9 DIV using an adapted calcium phosphate transfection protocol (12), as previously described (10).

### Electrophysiology

Voltage-clamp whole-cell recordings were performed stimulating Schaffer collateral fibers and recording evoked synaptic responses from CA1 pyramidal neurons at different holding potentials. The AMPA/NMDA ratios were calculated by acquiring AMPA receptor responses at –60 mV and NMDA receptor responses at +40 mV at a latency at which AMPAR responses were fully decayed (60 ms after stimulation). Picrotoxin (100 μM) was present in the external solution to block the GABA_A_ receptor responses. The recording chamber was perfused with external solution (in mM: 119 NaCl, 2.5 KCl, 1 NaH_2_PO_4_, 11 glucose, 26 NaHCO_3_, 4 MgCl_2_, 4 CaCl_2_ and 0.004 2-chloroadenosine, at pH 7.4), and was gassed with 5% CO_2_/ 95% O_2_. Patch recording pipettes (3–6 M Ω) were filled with internal solution (in mM: 115 CsMeSO_3_, 20 CsCl, 10 HEPES, 2.5 MgCl2, 4 Na_2_ATP, 0.4 Na_3_GTP, 10 sodium phosphocreatine and 0.6 EGTA, at pH 7.25). Synaptic responses were evoked with bipolar electrodes using single-voltage pulses (200 μs, up to 20 V). The stimulating electrodes were placed over Schaffer collateral fibers between 300 and 500 μm from the CA1 recorded cells. Synaptic responses were averaged over 50 trials. Whole-cell recordings were carried out with a Multiclamp 700A amplifier (Molecular Devices, Sunnyvale, USA).

### chemical LTP (cLTP) protocol

cLTP was induced as previously described (13). 19 DIV hippocampal cultures were washed with extracellular solution (ECS) containing (in mM): 150 NaCl, 2 CaCl_2_, 5 KCl, 10 HEPES, 30 Glucose, 0.001 TTX, 0.01 strychnine, 0.03 picrotoxin, pH 7.4. After washing, neurons were stimulated with glycine (300 μM) at room temperature for 3 min in ECS and then incubated for 20–25 min in ECS in a 37°C, 5% CO_2_/95% air incubator. Surface GluA1-AMPAR were labeled, and the cells were fixed and probed as previously described (10).

### Quantum dots labeling and Imaging

Endogenous GluA2 and GluA1-SEP labeling was performed in two steps: first neurons were incubated for 10 min at 37°C with anti-GFP antibody (1/300000) or anti-GluA2 antibody (1/1000), diluted in conditioned medium. After one washing step, anti-mouse IgG conjugated QD655 (diluted 1:10 in PBS) were diluted in conditioned medium with BSA 2% (1/2000) and were incubated on cells for 5 min at 37°C. Synapses were labeled using transfection with Homer1C-DsRed or Homer1C-GFP. All washes were performed in ECS containing (in mM) NaCl 145, KCl 5, Glucose 10, Hepes 10, CaCl_2_ 2 and MgCl_2_ 2), supplemented with BSA 2% at 37°C. After washing, neurons were mounted in an open chamber (K.F. Technology SRL) and imaged in ECS. Single-particle tracking was performed as in (14). Cells were imaged at 37°C on an inverted microscope (AxioObserver Z1, Carl Zeiss) equipped with a Plan Apochromat 63× oil objective (NA = 1.4). QDs, Homer1C-DsRed and Homer1C-GFP signals were detected by using a HXP fluorescence lamp (For QDs: excitation filter 425/50 and emission filters 655/30, Chroma). Fluorescent images from QDs were obtained with an integration time of 50 ms with up to 600 consecutive frames. Signals were recorded with a digital CMOS camera (ORCA Flash 4.0, Hamamatsu). QD-labeled GluAs were imaged on randomly selected dendritic regions over up to 30 min total experimental time. QDs fixed on the coverslip allowed us to compensate mechanical drifts of the stage.

### Quantum dots Tracking and Analysis

The tracking of single QDs was performed with homemade software based on Matlab (Mathworks Inc., Natick, USA). Single QDs were identified by their diffraction limited signals and their blinking fluorescent emission. The trajectory of a QD tagged receptor could not be tracked continuously due to the random blinking events of the QDs. When the positions before and after the dark period were compatible with borders set for maximal position changes between consecutive frames and blinking rates, the subtrajectories of the same receptor were reconnected. The values were determined empirically: 2–3 pixels (0.32–0.48 μm) for maximal position change between two frames and maximal dark periods of 25 frames (1.25 s). MSD curves were calculated for reconnected trajectories of at least 20 frames. The QDs were considered synaptic if colocalized with Homer dendritic clusters for at least five frames. Diffusion coefficients were calculated by a linear fit of the first 4–8 points of the mean square displacement (MSD) plots versus time depending on the length of the trajectory within a certain compartment. The resolution limit for diffusion was 0.0075 μm^2^/s as determined by(15), whereas the resolution precision was ∼40 nm. Statistical values are given as mean ± SEM or medians ± 25%/75% interval, if not stated otherwise. Statistical significances were performed using GraphPad Prism software. Non-Gaussian distributed data sets were tested by Mann-Whitney *t*-test. Indications of significance correspond to p values < 0.05 (^∗^), p<0.01 (^∗∗^), and p <0.001 (^∗∗∗^).

### Biochemistry

Protein extracts were prepared in lysis buffer [10 mM HEPES (pH 7.4), 150 mM NaCl, 10 mM EDTA, 1% (v/v) Triton X-100 supplemented with 1 mM DTT, 0.1 mM phenylmethylsulfonyl (PMSF), 1 μg/ml chymostatin, 1 μg/ml leupeptin, 1 μg/ml antipain, 1 μg/ml pepstatin (CLAP) and a cocktail of phosphatase inhibitors (1x, Roche, Carnaxide, Portugal)]. After centrifugation at 16,100 x *g* for 10 min at 4°C, protein in the supernatant was quantified using the bicinchoninic acid (BCA) assay kit (Pierce, Termo Fisher Scientific, Rockford, USA), and the samples were denatured with 5x concentrated denaturating buffer [62.5 mM Tris-HCl (pH 6.8), 10% (v/v) Glicerol, 2% (v/v) SDS, 0.01% (w/v) bromophenol blue and 5% (v/v) β-mercaptoethanol (added fresh)], and boiled for 5 min. Protein extracts were resolved by SDS-PAGE in 7.5% or 12% polyacrylamide gels. For western blot analysis, proteins were transferred onto a PVDF membrane (Millipore, Madrid, Spain) by electroblotting (40 V, overnight at 4°C). The membranes were blocked for 1 h at room temperature in Tris-buffered saline (137 mM NaCl, 20 mM Tris-HCl, pH 7.6) containing 0.1% (v/v) Tween-20 (TBS-T), and 5% (w/v) low-fat milk or BSA. Membranes were probed during 1 h, at room temperature, or overnight, at 4°C, with the primary antibodies diluted in TBS-T containing 5% or 0.5% (w/v) low-fat milk or 5% (w/v) BSA. Following several washes, membranes were incubated for 1 h with alkaline phosphatase-conjugated secondary antibodies (anti-mouse or anti-rabbit, depending on the primary antibody host species) at room temperature, washed again and incubated with chemifluorescent substrate (ECF) (GE Heathcare, Carnaxide, Portugal) for 5 min at room temperature. Membranes were scanned with the Storm 860 scanner (GE Heathcare, Carnaxide, Portugal), and quantified using the ImageQuant software under linear exposure conditions. When necessary, the membranes were stripped (0.2 M NaOH for 5 min) and re-probed.

